# Discovery of a Novel Non-Narcotic Analgesic Derived from the CL-20 Explosive: Synthesis, Pharmacology and Target Identification of Thio-wurtzine, a Potent Inhibitor of the Opioid Receptors and the Voltage-Dependent Calcium Channels

**DOI:** 10.1101/2021.03.18.435981

**Authors:** Stephanie Aguero, Simon Megy, Mailys Fournier, Daria A. Kulagina, Sergey V. Sysolyatin, Alexander I. Kalashnikov, Svetlana G. Krylova, Alexander B. Vorozhtsov, Vadim V. Zhdanov, Raphael Terreux

**Affiliations:** Équipe ECMO, Laboratoire de Biologie Tissulaire et d’Ingénierie (LBTI), UMR5305, Université Lyon 1, Lyon, France; Institute for Problems of Chemical and Energetic Technologies, Siberian Branch of the Russian Academy of Sciences (IPCET SB RAS), Biysk 659322, Altai Krai, Russia; Goldberg Research Institute of Pharmacology and Regenerative Medicine, Tomsk National Research Medical Center of the Russian Academy of Sciences, Tomsk 634028, Russia; National Research Tomsk State University, Lenin Avenue, 36, Tomsk 634050, Russia

**Keywords:** thiowurtzine, hexaazaisowurtzitane, analgesic activity, chemical synthesis, pharmacophore, pharmacokinetics, calcium channel, opioid receptor, docking, homology modeling, molecular dynamics

## Abstract

The number of candidate molecules for new non-narcotic analgesics is extremely limited. Here we report the identification of thiowurtzine, a new potent analgesic molecule with promising application in chronic pain treatment. We describe the chemical synthesis of this unique compound derived from the hexaazaisowurtzitane (CL-20) explosive molecule. Then we use animal experiments to assess its analgesic activity *in vivo* upon chemical, thermal and mechanical exposures, compared to the effect of several reference drugs. Finally, we investigate the potential receptors of thiowurtzine in order to better understand its complex mechanism of action. We use docking, molecular modeling and molecular dynamics simulations to identify and characterize the potential targets of the drug and confirm the results of the animal experiments. Our findings finally indicate that thiowurtzine may have a complex mechanism of action, by targeting the mu, kappa, delta and ORL1 opioid receptors, and the voltage-gated calcium channels as well.

## INTRODUCTION

Synthesis of candidate molecules for the design of non-narcotic analgesics to relieve severe and moderate pain is a trending topic in pharmaceutics. Three types of treatments against pain of different etiologies are primarily used in out-patient and in-patient: (1) opioid and cannabinoid molecules with various analgesic activity, (2) nonsteroidal anti-inflammatory drugs (NSAIDs) acting as antalgic molecules, and (3) their various combinations in multimodal approaches for chronic pain, incorporating modalities such as physiotherapy, psychological therapy, patient education and peripheral stimulation.^1 2 3 4 5^ An alternative to opiates for severe and moderate pain relief may reside in the use of non-narcotic analgesics acting on the central and peripheral nervous systems. However, the potential candidate molecules for non-narcotic analgesics are extremely limited in number. In this regard, one of the current successful approaches for the discovery of new molecules for biomedical research is the development of high-throughput methods of virtual screening for large databases of chemical compounds. ^6 7 8^

Here we report the development of a new patented anagesic molecule, ^9^ hereafter referred as thiowurtzine, and based on the hexaazaisowurtzitane (CL-20) molecule, a substance commonly known as a component of rocket fuel and explosives. Although several explosives such as nitroglycerin or potassium nitrate have been used as drugs in the medical history, ^10 11 12 13^ the case of thiowurtzine is quite different, as it is directly derived from the CL-20 molecule, but lacks all of the 6 nitro groups which confer its explosive properties to CL-20. As the 6 nitro groups are only added during the very last step of the chemical synthesis of CL-20, ^14^ thiowurtzine can very well be considered more as a modified precursor rather than a revised explosive molecule.

Hexaazaisowurtzitane derivatives are caged polycyclic polynitrogen compounds and possess a framework structure determining their unique properties. ^15 16^ Yet, numerous studies focused on the synthesis of new hexaazaisowurtzitane derivatives have until recently been primarily focused on their explosive and propelling properties for military purposes, as promising components of solid rocket fuels and composite explosives. ^16 17^

Only in recent years has hexaazaisowurtzitane been investigated as a new pharmacophore for the development of original pharmaceutical substances. In this more biological context, preliminary studies investigating the analgesic activity of hexaazaisowurtzitane derivatives have recently been described: lately, the analgesic activity of thiowurtzine has been characterized in mice. ^18^ Its pharmacokinetics in rats ^19^ and its concomitant use with cyclophosphamide in mice cancer therapies ^20^ have been investigated as well. The aim of the project was to conduct conclusive preclinical studies of this drug based on the hexaazaizowurtzitane molecule for the treatment of pain syndrome caused by different etiologies.

Thiowurtzine is the first chemical compound in the class of hexaazaisowurtzitanes which presents a significant, experimentally validated bioactivity. ^21 15^

In this study, we report the synthesis of 4-(3,4-dibromothiophenecarbonyl)-2,6,8,12-tetraacetyl-2,4,6,8,10,12-hexaazatetracyclo[5,5,0,0^3,11^,0^5,9^] dodecane (thiowurtzine, **I**), whose properties were predicted using the PASS software package. ^22^ Furthermore, we use animal experiments to assess the analgesic activity of thiowurtzine *in vivo* upon chemical, thermal and mechanical exposures, compared to the effect of several reference molecules. Finally, we investigate the potential receptors of thiowurtzine in order to better understand its mechanism of action. We use docking, molecular modeling and molecular dynamics simulations to identify and characterize the potential targets of the drug and confirm the results of the animal experiments.

## RESULTS AND DISCUSSION

### Chemistry

A series of sequential reactions were used for the synthesis of thiowurtzine (I) as a final compound, as illustrated in Scheme 1.

**Scheme 1.**
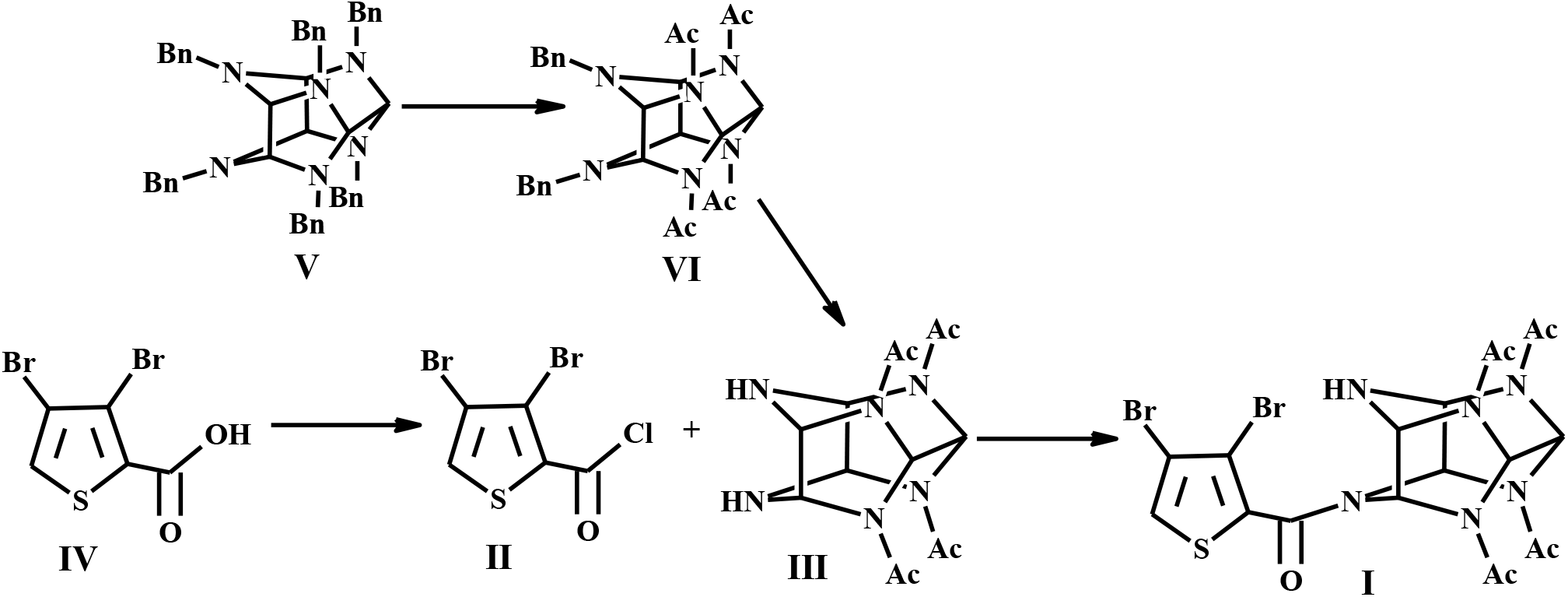
Synthesis of thiowurtzine (I)

For the synthesis of compound **IV**, we optimized the procedure involving exhaustive bromination of thiophene, reduction with zinc dust to dibromothiophene, alkylation in the presence of titanium tetrachloride, and final oxidation of the aldehyde to the desired dibromothiophene carboxylic acid **IV**.

The yield of **IV** on a thiophene **(VII)** basis was 37%. In case compound **VIII** was used, the yield of **IV** was 73%. These reactions are illustrated in Scheme 2. As pictured in Scheme 1, acylating agent **II** was obtained in a quantitative yield by treating 3,4-dibromothiophene carboxylic acid **IV** with boiling thionyl chloride for 1 hour.

**Scheme 2.**
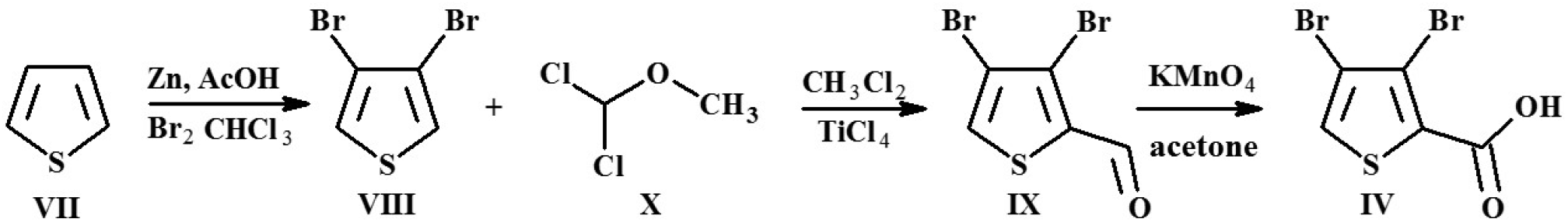
An optimized procedure for dibromothiophene carboxylic acid (IV)

Synthetic methods for tetraacetyl hexaazaisowurtzitane (**III**) are commonly known and industrially employed.^23 24 25^ A cascade condensation of benzylamine with glyoxal furnished hexabenzyl hexaazaisowurtzitane (**V**). A twostage hydrogenation of **V** over Pd catalyst produced **VI** (Scheme 1).

The acylation reaction between the starting **III** and chloroanhydride **II** was performed in boiling acetonitrile. The use of other organic solvents diminished the yield of product **I**. To lower the quantity of impurities, diamine **III** in acetonitrile was treated with the acylating agent **II**, with compound **I** precipitating from the reaction mixture and not undergoing further reactions (Scheme 1).

No excess acylating agent **II** was required to complete the reaction. When **II** was used in excess from 0% to 65%, the yield of **I** ranged from 79% to 82%.

The structure of the obtained compound **I** was verified by a complex approach via physicochemical methods, including ^1^H and ^13^C NMR spectra (as illustrated in the Supporting Information).

### Biological Evaluation

The predicted analgesic activity of the resulting compound **I** was confirmed on animals using the reference drugs tramadol, ketorolac, and diclofenac.

Two experimental series with chemical pain stimulation of the peritoneum showed a pronounced antinociceptive effect of thiowurtzine upon a single (100 mg/kg) and sub-chronic intragastric administration to mice in doses of 50 and 100 mg/kg (Table 1). After a single administration (100 mg/kg), thiowurtzine significantly relieved acute visceral pain by 56.4% and prolonged the pain response time (p<0.05) in reference to the negative control and appeared to be equivalent to the reference drug tramadol, which exhibits a 62.2% pain response inhibition (Table 1, series I). It should be noted that in the second series of experiments, after a 4-day compound administration in doses of 50 and 100 mg/kg, thiowurtzine reduced the number of writhings by 56.9% (p<0.01) and 47.1% (p<0.01) in comparison with the control, respectively (Table 1, series II). Thiowurtzine (50 mg/kg) exceeded in activity the reference drug diclofenac.

**Table 1.** Anti-nociceptive effect of thiowurtzine in acetic-acid writhing test (X±m)

| Group | Number of writhings within 20 min | Writhing latency, sec | Pain response inhibition, % |
| --- | --- | --- | --- |
| Series I: inbred male CBA mice. Single administration of thiowurtzine and Tramadol |  |  |  |
| 1. Negative control — vehicle (n=9) | 24.1±3.4 | 160.9±86.4 | 0 |
| 3. Tramadol, 20 mg/kg (n=9) | 9.1±2.0<br>** (1-3) | 444.8±42.8<br>** (1-3) | 62.2 |
| 5. Thiowurtzine, 100 mg/kg (n=6) | 10.5±3.0<br>* (1-5) | 382.5±120.4<br>* (1-5) | 56.4 |
| Series II: outbred female mice, stock CD1. Drug administration for 4 days |  |  |  |
| 1. Negative control — vehicle (n=10) | 27.4±3.1 | 276±15 | - |
| 2. Diclofenac, 10 mg/kg (n=10) | 18.2±1.2<br>** (1-2) | 270±10 | 33.6 |
| 3. Thiowurtzine, 50 mg/kg (n=10) | 11.8±2.9<br>** (1-3)<br>* (2-3) | 270±10 | 56.9 |
| 4. Thiowurtzine, 100 mg/kg (n=10) | 14.5±3.0<br>** (1-4) | 273±16 | 47.1 |

**Note.** \* $p < 0.05$ , \*\* $p < 0.01$ compared to the negative control and reference group (Mann—Whitney U test).

The most pronounced analgesic effect of thiowurtzine was revealed in the central algesia model, which is controlled by the cortical and subcortical structures of brain. In the hot plate test, subchronic intragastric administration of thiowurtzine in a dose of 100 mg/kg caused a maximum increase in the pain response latency 77.9% — by 1.8 times (p<0.01) relative to the control (Table 2, series I, II). Thiowurtzine (100 mg/kg) exceeded the activity of the reference drug diclofenac, which pain response inhibition proved to be 11.3%.

**Table 2.** Analgesic activity of thiowurtzine in hot-plate test in outbred male stock CD rats (X±m)

| Group | Pain response latency, sec | Pain response inhibition, % |
| --- | --- | --- |
| Series I: Drug administration for 4 days |  |  |
| 1. Negative control — vehicle (n=8) | 9.8±1.0 | - |
| 2. Diclofenac, 5 mg/kg (n=9) | 10.9±1.5 | 11.3 |
| 3. Thiowurtzine, 25 mg/kg (n=9) | 10.4±1.3 | 6.1 |
| 4. Thiowurtzine, 50 mg/kg (n=8) | 12.1±1.6 | 23.5 |
| Series II: Drug administration for 4 days |  |  |
| 1. Negative control — vehicle (n=10) | 11.3±1,0 | - |
| 2. Thiowurtzine, 100 mg/kg (n=10) | 20.1 ± 2.3<br>**(1-2) | 77.9 |
| 3. Thiowurtzine, 200 mg/kg (n=10) | 14.9±2.1<br>*(2-3) | 31.9 |
| 4. Thiowurtzine, 300 mg/kg (n=10) | 13.2±2,7 | 16.8 |

**Note.** \*p<0.05, \*\*p<0.01 compared to the negative control and reference group (Mann—Whitney U test).

Thiowurtzine showed a roughly constant increase in the nociceptive threshold throughout the study, which was noted to be significant in the 100 mg/kg group (Table 3). After a single administration (100 mg/kg), thiowurtzine significantly increased the supporting pressure in the left hind paw pad by 2.1 times (1 h, p<0.01) and 1.6 times (4 h, p<0.01), relative to the negative control.

**Table 3.** Analgesic activity of thiowurtzine in Randall-Selitto test in outbred male stock CD1 mice (X±m)

| Group | Supporting pressure in left hind paw pad, g/ Condition |  | Supporting pressure in right hind paw pad with formalin edema, g/ Condition |  |
| --- | --- | --- | --- | --- |
|  | 1 hour after drug administration | 4 hours after drug administration | 1 hour after drug administration<br>1-5 min after injection of formalin | 1 hour after drug administration<br>40-50 min after injection of formalin |
| 1. Negative control — vehicle (n=10) | 294.9±48.4 | 373.2±70.9 | 141.1±23,7 | 305.2±72.6 |
| 2. Tramadol, 10 mg/kg (n=9) | 645.2±40.8<br>**(1-2) | 606.7±53.9<br>**(1-2) | 318.9±84.1<br>*(1-2) | 518.0±85.0<br>*(1-2) |
| 3. Ketorolac, 6 mg/kg (n=8) | 521.2±107.9<br>**(1-3) | 449.7±90.4 | 491.4±88.2<br>*(1-3) | 396.6±94.1 |
| 4.Thiowurtzine, 100 mg/kg (n=9) | 631.5±43.9<br>**(1-4) | 612.9±43.8<br>**(1-4) | 620.8±60.0<br>**(1-4) | 611.3±58.8<br>**(1-4) |
| 5. Thiowurtzine, 200 mg/kg (n=9) | 586.1±68.6<br>**(1-5) | 455.8±81.7 | 419.7±94.2<br>*(1-5) | 366.3±88.3 |

**Note.** \*p<0.05, \*\*p<0.01 compared to the negative control and reference group (Mann—Whitney U test).

Interestingly, the Randall-Selitto test performed on the right paw (100 mg/kg) with formalin edema also revealed a significant increase in the nociceptive threshold. In this case, an increased supporting pressure was evidenced, by 4.4 times (1 h, p<0.01) and 2.0 times (4 h, p<0.01) compared to the negative control.

The assessed effect of thiowurtzine (100 mg/kg) on thresholds of response to mechanical pressure stimulation appeared to be comparable with tramadol but exceeded the activity of the reference drug ketorolac.

The preclinical study demonstrated that chronic administration of thiowurtzine (abstinence syndrome) did not evoke drug abuse. There was no impact on the respiration and the central nervous systems, indicating that thiowurtzine is not a morphine-like action compound, and does not cause ulcerogenic damage to the gastrointestinal mucosa of the test animals.

### Target Identification

Preliminary studies investigating the analgesic activity of thiowurtzine in conjunction with naloxone and naloxone methiodide have been previously published.^18^ Naloxone has been shown to have a higher affinity than naloxone methiodide for opioid mu, kappa, and delta receptors in the brain of the mouse. ^26^ Therefore naloxone targets the brain tissue, whereas naloxone methiodide, which has limited access to the brain, targets more specifically the peripheral opioid receptors.

In particular, injection of the non-selective opioid receptor antagonist naloxone was shown to weaken the analgesia caused by a single thiowurtzine administration in a dose of 100 mg/kg, suggesting an action on the opioid mu, kappa, and delta receptors in the central opioid system of the mouse.^18^

On the contrary, naloxone methiodide did not abolish analgesia, suggesting that thiowurtzine has no affinity for the peripheral opioid receptors.

These results are consistent with the central opioid system being involved in the mediation of the analgesic effect of thiowurtzine. However, the antinociceptive effect of thiowurtzine was not completely abolished by naloxone, which suggests a complex mechanism of action of the compound under study. In these regards, the opioid analgesia pathway seems to be only one component of the mechanism of action of thiowurtzine, and consequently there may be other receptors targeted.

In order to identify new potential targets for the action of thiowurtzine, we screened the Drugbank^27^ database looking for structurally similar drugs and their well-known receptors. In particular, we identified several drugs classified as calcium antagonists and acting as calcium channel blockers (namely pranidipine, nicardipine, and especially lacidipine, which exhibited the highest score of 86% for structural similarity with thiowurtzine). Lacidipine, like other dihydropyridines, targets the voltage dependent calcium channels, and has a selective activity on the Ca_v_ 1.2. channel. Dihydropyridines have a high affinity for these receptors ranging from 0.1 to 50 nM, and they specifically interact with their α_1c_ subunit.^28^ This led us to investigate this class of voltage-gated calcium channels as potential targets for thiowurtzine as well. Indeed, calcium channel blockers have the ability to inhibit voltage-gated calcium channels and thus to reduce the release of neurotransmitters at the presynaptic level, which may lead to pain reduction. As a consequence, voltage-gated calcium channels have been described as targets of choice for potential analgesics in the previous years^29 30 31 32 33^, especially when used concomitantly with opioids in attenuation of clinical pain.^34 35 36^ In this particular case, calcium channel blockers have been shown to increase morphine analgesia.^37 38^

We used the GOLD^39^ (Genetic Optimization for Ligand Docking) software to perform docking experiments of the thiowurtzine molecule on the mu, kappa, and delta opioid receptors, the opioid like receptor 1 (ORL1), and on the voltage dependent calcium channel Ca_v_ 1.2 as well

The mu, kappa, delta and ORL1 opioid receptors were constructed using the corresponding UniProtKB human sequences and the I-TASSER software (Iterative Threading ASSEmbly Refinement) from the Zhang Lab. ^40^ The models were then checked, charged, and minimized with the MOE software using the AMBER 14:ETH force field.

As no satisfactory structure was available for the mice calcium channel Ca_v_ 1.2 interacting with a dihydropyridine, we also built a homology model using the UniProtKB sequence Q13936 (isoform 1) of the human voltage-gated calcium channel subunit α_1c_ Ca_v_ 1.2 and the PDB structure 6JP5 of the rabbit Ca_v_1.1 / nifedipine complex as a template. The RMSD calculated between the final model and the template was 0.738 Å. The RMSD between the binding sites (residues with atoms within a sphere of 4.5 Å around the nifedipine ligand) of 6JP5 and the homology model was 0.32 Å, which indicates that the two binding sites are extremely similar. The Ramachandran diagram of the model was also checked with 93% of residues in the most favorable regions, 6% in the allowed regions, and only a few outlier residues. The homology model was further evaluated by docking 9 ligands extracted from the BindingDB (http://bindingdb.org). The docking scores were in very good agreement with the experimentally measured inhibitor constants (Ki) for each of the 9 ligands, thus validating the robustness of our model.

### Docking of Thiowurtzine

The docking of thiowurtzine into the models of the five receptors (mu, kappa, delta, ORL1, and Ca_v_ 1.2) was performed using the GOLD software. We used control drugs (tramadol, ketorolac, and diclofenac) as references. As tramadol is commercially available as a racemic mix (1*R*-2*R* and 1*S*-2*S*), we separately tested each of the two enantiomers and reported their respective score. All the results are summarized in Table 4.

**Table 4.** Docking scores of thiowurtzine and control drugs against mu, kappa, delta, ORL1, and Ca_v_ 1.2 receptors.

| Drug | GOLD score with the mu receptor | GOLD score with the kappa receptor | GOLD score with the delta receptor | GOLD score with the ORL1 receptor | GOLD score with the Ca<sub>v</sub> 1.2 receptor |
| --- | --- | --- | --- | --- | --- |
| Tramadol 1*S*-2*S* | 19,546 | 27,003 | 21,797 | 20,520 | 24,499 |
| Tramadol 1*R*-2*R* | 19,188 | 28,349 | 20,435 | 19,272 | 23,597 |
| Ketorolac | 19,529 | 26,150 | 23,640 | 19,467 | 22,065 |
| Diclofenac | 20,823 | 31,603 | 24,274 | 17,567 | 24,154 |
| Thiowurtzine | 8,465 | 2,730 | 6,407 | 2,153 | 15,562 |

Note. The scores of the GOLD<sup>39</sup> docking software using the Chemscore<sup>41 42</sup> rescore function are displayed.

Interestingly, the docking scores of thiowurtzine are lower than those of the reference drugs for all the investigated receptors. However, these scores are high enough to unequivocally demonstrate an effective binding of thiowurtzine, especially with the mu, delta and Ca_v_ receptors. For the mu and delta receptors, the scores are about 2.5 to 4 times lower than those of tramadol, ketorolac or diclofenac. However, the animal experiments evidenced that the doses of thiowurtzine needed to be typically 5 times higher than those of tramadol (20 mg/kg vs 100 mg/kg) and diclofenac (10 mg/kg vs 50 mg/kg) to produce similar effects with the acetic acid writhing test (Table 1). The same ratio was also observed for the hot plate test (diclofenac 5 mg/kg, thiowurtzine 25 mg/kg, Table 2) to produce similar results. Aside from the typical bioavailability issues, the animal experiments are thus in excellent agreement with our docking results. The best docking pose for the thiowurtzine in complex with the model of the mu opioid receptor is displayed in Figure 1.

**Figure 1.**
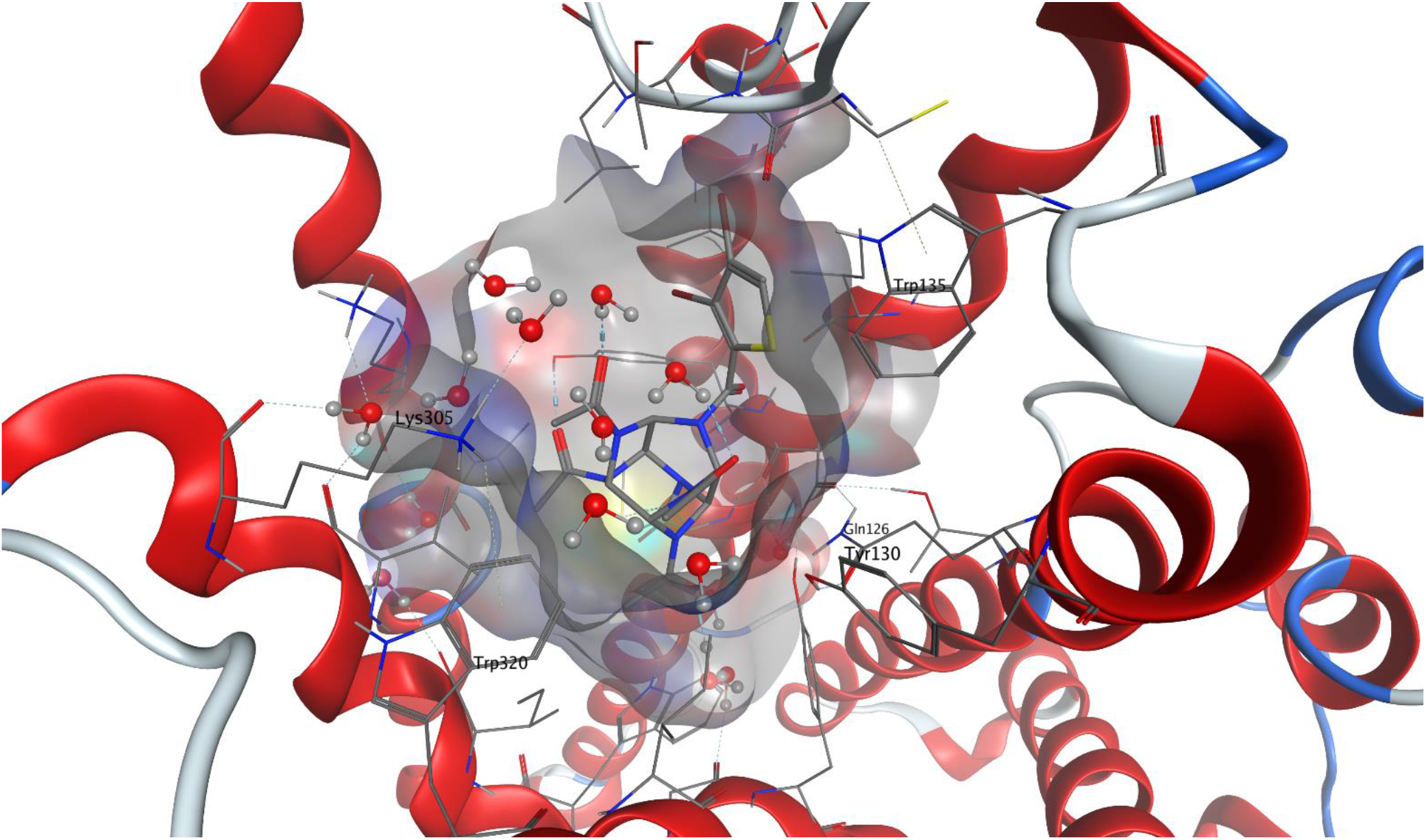
Molecular representation of the best docking pose of the thiowurtzine molecule interacting with the calculated model of the mu opioid receptor. Thiowurtzine is displayed as sticks, and the mu receptor with red ribbons. The water molecules are visible, as well as the molecular surface of the cavity surrounding the thiowurtzine molecule. The sidechains of the residues interacting with thiowurtzine are labelled and displayed as sticks as well.

For the Ca_v_ receptor, the docking scores of thiowurtzine are much better than for any of the other investigated receptors (Table 4), and they are close to those of the other reference drugs (albeit being about 30% lower). Consequently, thiowurtzine appears to be a significantly better ligand for the Ca_v_ receptor than for any of the investigated opioid receptors. This is consistent with a complex mechanism of action, which could target at the same time the opioid receptors and the voltage-gated calcium channels.

### Molecular Dynamics

In order to validate whether our hypothesis of thiowurtzine binding to the Ca_v_ 1.2 calcium channel was stable over time, molecular dynamics experiments were carried out for the receptor in presence of thiowurtzine. We used the best docking pose of thiowurtzine with the model of the Ca_v_ 1.2 calcium channel as a starting point. A lipid bilayer surrounding the calcium channel was generated with the MOE lipid generator. Water molecules were added, as well as Na+ and Clions in order to simulate a 0.1 M ionic force, and the pH was set at 7.4. Standard Amber 14:EHT parameters were used for the molecular dynamics (MD). A quick minimization was performed prior to the start of the MD simulation. After a preliminary equilibration phase, a 200 ns molecular simulation was performed. A frame was saved every 200 ps, leading to a total number of 1,000 frames. The last frame of the simulation is displayed in Figure 2.

**Figure 2.**
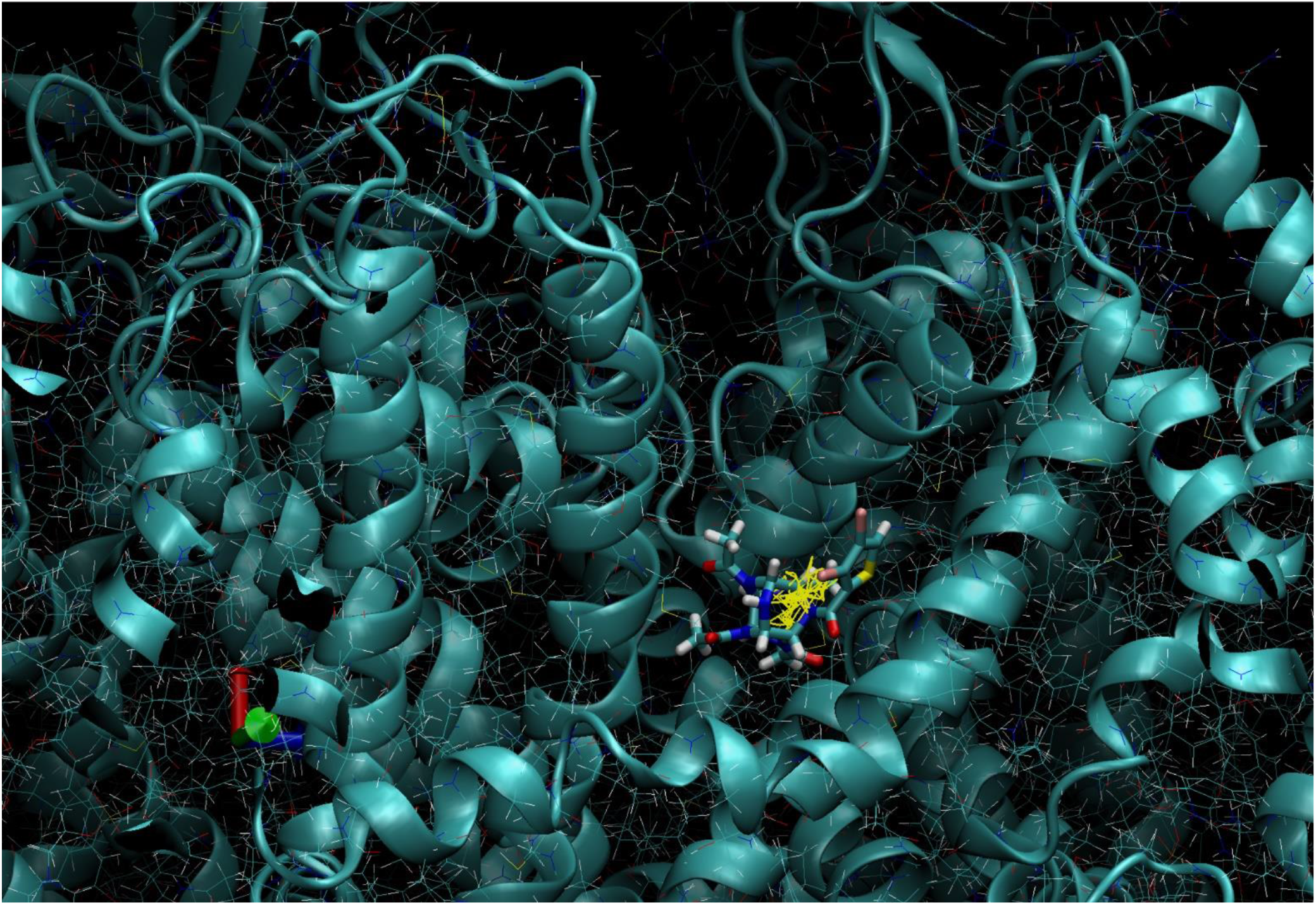
Molecular representation of the last frame of the 200 ns molecular dynamic simulation of thiowurtzine in complex with the model of the Ca_v_ receptor. Thiowurtzine is displayed as sticks, and the Ca_v_ calcium channel with teal ribbons. The trajectory of the thiowurtzine mass center throughout the entire simulation is displayed with yellow lines.

The molecular complex formed by thiowurtzine and the Ca_v_ calcium channel model appeared to be extremely stable over time. In fact, thiowurtzine stayed within the interaction site during the 200 ns of the simulation, with only limited relative motions around its starting position at the beginning of the simulation. The full trajectory of the thio-wurtzine mass center during the complete simulation is displayed with yellow sticks in Figure 2 and evidences a great stability of the complex over time. These results are in excellent agreement with the good docking scores previously obtained with the GOLD software for the thio-wurtzine interacting with the model of the Ca_v_ receptor (Table 4).

## CONCLUSION

In summary, we provided the chemical synthesis, the biological evaluation and the *in-silico* target identification of a potent new hexazaisowurtzitane-based analgesic referred here as thiowurtzine. This non-addictive analgesic does not exhibit any of the NSAIDs typical adverse effects and may very well lead to new compounds of the same class with practical applications.

It is worth noting that the devised original method for the synthesis of 3,4-dibromothiophene carboxylic acid holds promise for further use, as it provides a quite high yield under mild synthesis conditions.

The experimental evidence (behavioral tests with thermal and chemical stimuli, and mechanical compression) demonstrates an analgesic effect of thiowurtzine through its inhibitory action on the peripheral and central mechanisms of development and maintenance of pain syndrome. These findings highlight a promising potential of the new compound as an analgesic against pain syndromes of different etiologies.

The software-based evaluation of thiowurtzine, which was validated by the animal studies, gave some interesting perspectives about the potential molecular receptors of thiowurtzine. We highlighted the role of the opioid receptors as primary targets, which confirmed the results of the experiments using thiowurtzine in conjunction with naloxone. However, the antinociceptive effect of thiowurtzine was not completely abolished by naloxone^18^, which suggests that the opioid analgesia pathway may not be the only component of the mechanism of action of thiowurtzine, and consequently, that there may be other receptors targeted as well.

Molecular comparisons with other drugs, associated with molecular modeling and molecular dynamics simulations allowed us to also consider voltage-gated calcium channels as potential targets. Interestingly, the possibility that calcium channel blockers might induce analgesia by reducing the release of neurotransmitters and thus increase morphine analgesia has been investigated and confirmed in several studies. ^34 35 36 37 38^ Those results are consistent with thiowurtzine having a primary action on the central opioid system and suggest that this primary action may very well be potentiated by a secondary action on the voltage-gated calcium channels as well. These results were confirmed by the docking scores of thiowurtzine against the opioid and the calcium receptors. Those findings might provide a solid basis for the explanation of the complicated mechanism of action of thiowurtzine.

New rational design studies involving further chemical modifications of hexaazaizowurtzitane-derivatives are currently under way and might lead to the discovery of a whole new class of biologically active molecules of interest.

## EXPERIMENTAL SECTION

### General Methods

High-performance liquid chromatography (HPLC) was performed on an Agilent Technologies 1260 Infinity instrument with a 2.1×15-mm pre-column, Zorbax SB-18 sorbent, and 3 μm fractions; 3.0×150-mm column, Zorbax SB-18 sorbent, 3.5 μm fractions using gradient elution. Eluent A: 0.2% orthophosphoric acid solution, eluent B: acetonitrile. ^1^H and ^13^C NMR spectra were recorded on a Bruker AM-400 spectrometer operating at 400.13 MHz for ^1^H and at 100.61 MHz for ^13^C; DMSO-*d*_*6*_ was used as solvent. IR spectra were taken on a Infralum FT-801 spectrophotometer in KBr pellets (1 mg substance in 200 mg potassium bromide) in a range from 4000 to 400 cm^−1^. The melting point was measured on a Stuart SMP 30 melting point apparatus.

### Synthesis of 2,4,6,8,10,12-hexabenzyl-2,4,6,8,10,12-hexaazatetracyclo[5,5,0,0^3,11^,0^5,9^]dodecane (V)

Into a stirred flask was put benzylamine (170 mL, 1.56 mol), distilled water (130 mL), acetonitrile (1430 mL) and 98% formic acid (5.4 mL). Then, 40% aqueous glyoxal (94.25 g, 0.65 mol) was added portionwise for 1 h at a temperature not above 20°C. The reaction mixture was held at room temperature for 17 h. The resultant crystalline product was collected by filtration and washed in-situ with cold acetonitrile. Yield: 121 g (76% on a benzylamine basis). Mp: 145–150°C.

For deresination, the resultant crude product was stirred in acetonitrile (240–250 mL) at 50°C for 15–20 min, cooled to room temperature, filtered and washed with acetonitrile to furnish 115–118 g of the product (Mp: 150–153°C) after drying. Calcd for C_48_H_48_N_6_ (%): 81.32; H 6.82; N 11.85; found (%): C 82.13; H 6.79; N 11.09. IR (*v*/cm^−1^): 3082, 3061, 3023, 2923, 2857, 2823, 1493, 1452, 1351, 1323, 1302, 1244, 1140, 1072, 987, 898, 835, 733, 699, 625. ^1^H NMR (DMSO-*d*_*6*_, δ, ppm): 3.48 (s, 2H, CH), 4.01 (s, 4H, CH2), 4,04 (s, 4H, CH2), 4.15 (s, 4H, CH), 7.14-7.26 (m, 30H, CHap).

### Synthesis of 4,10-dibenzyl-2,6,8,12-tetraacetyl-2,4,6,8,10,12-hexaazatetracyclo[5,5,0,0^3,11^,0^5,9^]dodecane (VI)

Into a 300-mL autoclave equipped with an electromagnetic stirrer and a hydrogen gas delivery system was loaded hexabenzylhexaazaisowurtzitane (10 g), bromobenzene (0.18-0.20 mL), a catalyst, and DMF (40 mL). Lastly, acetic anhydride (15 mL) was added, and the autoclave was closed. The autoclave was purged with hydrogen three times. Hydrogen was then supplied under 5-6 kgf/cm^2^, stirring was turned on (550-600 rpm), and the whole was heated to 50-55°C. The hydrogen absorption was controlled against the autoclave pressure drop. If the autoclave pressure went down below 2.5–2 kgf/cm^2^, hydrogen was again fed to 5–6 kgf/cm^2^.

After the process was complete (an autoclave pressure drop of 0.1-0.2 kgf/cm^2^ within 1 h), the heating was turned off, cold water was poured into the bath, and the reactor was cooled to 20-25°C.

Hydrogen depressurization was then performed, and the product suspension with the catalyst was filtered. The mixture of dibenzyltetraacertylhexaazaisowurtzitane (DBTA) and catalyst was washed with ethanol (3 x 10 mL), squeezed and air-dried to furnish DBTA not allowing for the catalyst weight. Yield: 5.8–6 g (80-83%). Calcd for C_28_H_32_N_6_O_4_ (%): C 65.10; H 6.24; N 16.27; O 12.39; found (%): C 64.79; H 6.18; N 16.25. IR (*v*/cm^−1^): 3083, 3044, 3022, 3005, 2876, 2829, 1687,1650, 1455, 1412, 1358, 1319, 1300, 1239, 1147, 1068, 985, 893, 834, 730, 687, 627. ^1^H NMR (DMSO-*d*_*6*_, δ ppm): 1.85-1.95 (m, 12H, CH3), 4.09 (s, 4H, CH2), 5.71 (br.s, 4H, CH), 6.51 (br.s 4H, CH), 7.37-7.49 (m, 10H, CHap).

### Synthesis of 2,6,8,12-tetraacetyl-2,4,6,8,10,12-hexaazatetracyclo[5,5,0,0^3,11^,0^5,9^] dodecane (III)

Into an autoclave was loaded DBTA (25 g), 5 % Pd/C (5 g) and 50% acetic acid (150 mL). The autoclave was closed, purged 3 times with nitrogen and 2 times with hydrogen. Hydrogen was then fed at 5 kgf/cm^2^ and stirring was turned on. The reaction was carried out at 70–75°C for 3 h. The catalyst was then filtered off, and the reaction mixture was evaporated in a rotary evaporator until it was viscous. To the evaporated mixture was added alcohol (150–200 mL) and the whole was stirred for 15–20 min. The crystalline product was collected by filtration and air-dried to give tetraacetylhexaazaisowurtzitane (TA). Yield: 15 g (92%), purity 98% (as per HPLC). Mp: 360°C (decomposition). Calcd for C_14_H_20_N_6_O_8_ (%): H 49.99; N 5.99; N 24.99; O 19.03; found (%): C 50.16; H 6.04; N 24.72. (IR *v*/cm^−1^): 3369, 3329, 3047, 3019, 2931, 1659, 1402, 1359, 1293, 1255, 1140, 1111, 1019, 974, 909, 801, 710, 609, 509. ^1^H NMR (DMSO-*d*_*6*_, δ ppm): 1.94-2.07 (m, 12H, CH3CO), 4.59-4.99 (m, 2H, NH), 5.29 (br.s, 2H, NH), 5.99-6.23 (m, 4H, CH).

Synthesis of 3,4-dibromothiophene carboxylic acid (IV): Into a round-bottom flask was put thiophene (48 mL, 0.6 mol), chloroform (24 mL), and bromine (128 mL, 2.4 mol) was slowly added dropwise. The dosing time was 4 h. In the beginning of the dosing, the solution was observed to be decolored and hydrogen bromide to evolve vigorously. Hydrogen bromide was entrapped and absorbed with a 20% NaOH solution. After dosing was completed, the reaction mixture was cooled on ice bath and held for 12 h.

Upon completion of holding, to the reaction mixture was added chloroform (20 mL) and the whole was heated on silicone bath at 85°C, and a 15% KOH solution in ethanol was added. The mixture was refluxed for 3 h with vigorous stirring, then cooled down and poured out onto ice (400 g). After holding for 1 h, the product was filtered off, washed with water and air-dried.

To a 1-L flask fitted with a distiller was added zinc powder (180.6 g, 2.76 mol) in water (240 mL), and glacial acetic acid (180 mL, 5.2 mol) was then added. The reaction mixture was heated to boil, and the crude product was added in a few portions. The distilled organic fraction was washed with water and then dried over MgSO_4_ to yield 3,4-dibromothiophene as a colorless liquid. Yield: 74.07 g (51%).

To a mixture of the resultant 3,4-dibromothiophene (0.024 mol) and dichloromethyl methyl ether (2.98 g, 2.35 mL, 0.026 mol) in dry dichloromethane (15 mL) was added TiCl4 (7.59 g, 4.39 mL, 0.04 mol) with stirring at −12°C for 40-50 min. The reaction mixture was stirred for 1 h, and water (30 mL) was added drop by drop within 15 min. After 30 min, the layers were separated, and the water layer was extracted with dichloromethane (3×30 mL). The combined extract was dried over MgSO_4_ and evaporated. The product was recrystallized from heptane, dried and used in the subsequent stage.

The resultant 3,4-dibromo-2-thiophene-aldehyde (47,1 g, 0.174 mol) was dissolved in acetone (550 mL). A potassium permanganate (37.8 g) solution in distilled water (720 mL) was prepared separately by heating.

To the aldehyde acetone solution was added potassium permanganate in portions of 15-20 mL at not above 20°C. The whole was held for 3-5 min after adding each portion. The dosing was stopped after the solution turned a stable pink color. The resultant suspension was decolored by adding a 10% sodium sulfite solution. The reaction mixture was filtered off of MnO_2_and washed with water.

The filtrate was evaporated by half and acidified with HCl until pH was less than 2. The product was collected by filtration, washed with water and then recrystallized from dichloroethane to afford 3,4-dibromothiophene carboxylic acid. Yield: 42.29 g (85%). Mp: 205-207°C. Calcd for C_5_H_2_Br_2_O_2_S (%): C 21.00; H 0.70; Br 55.89; O 11.19; S 11.21l found: (%): C 21.09; H 0.69; S 11.24. IR (*v*/cm^−1^): 3102, 3098, 1663, 1478, 1389, 1321, 1123, 920, 857, 784, 685. ^1^H NMR (DMSO-*d*_*6*_, δ ppm): 7.44 (c, 1H, CH), 10.93 (br.s, 1H, OH).

### Synthesis of 3,4-dibromothiophenecarbonyl chloride (II)

3,4-dibromothiophene carboxylic acid (10 g, 0.035 mol) was put into thionyl chloride (50 mL) and heated to the boiling point of thionyl chloride. The mixture was held at the same temperature for 1 h, then cooled down and evaporated in a rotary evaporator. Yield: 9.2 g (86%). Mp was not measured because the product was immediately used in further reaction.

Synthesis of 4-(3,4-dibromothiophenecarbonyl)-2,6,8,12-tetraacetyl-2,4,6,8,10,12-hexaazatetracyclo[5,5,0,0^3,11^,0^5,9^]dodecane (I). To a solution of chloroanhydride V (9.2 g, 0.03 mol) in dry acetonitrile (60 mL) was added 2,6,8,12-tetraacetyl-2,4,6,8,10,12-hexaazatetracyclo[5,5,0,0^3,11^,0^5,9^]dodecane (5.04 g, 0.015 mol). The resultant mixture was refluxed for 2 h and then cooled down, and the precipitated product was collected by filtration and washed 2 times with acetonitrile.

The dried creamy-colored sediment was recrystallized from 70% (by vol.) aqueous ethanol. The resultant product was washed with water 3 times and air-dried to furnish 4-(3,4-dibromothiophenecarbonyl)-2,6,8,12-tetraacetyl-2,4,6,8,10,12-hexaazatetracyclo[5,5,0,0^3,11^,0^5,9^]dodecane as a colorless crystalline product with an assay of 99% (as per HPLC). Yield: 7.19 g (79.4%). MP: 332-334°C. Calcd for C_19_H_20_Br_2_N_6_O_5_S (%): C 37.76; H 3.34; Br 26.45; N 13.91; O 13.24; S 5.31; found (%): C 37.68; H 3.24; N 13.82; S 5.37. IR (*v*/cm^−1^): 3287, 3119, 3005, 1657, 1504, 1425, 1356, 1323, 1289, 1250, 1162, 1045, 984, 893, 878, 742, 715, 623. ^1^H NMR (DMSO-*d*_*6*_, δ ppm): 1.85-2.15 (m, 12H, CH3CO), 4.75-5.12 (m, 1H, NH), 5.45-5.67 (m, 2H, CH), 5.92-6.85 (m, 4H, CH), 8.08-8.18 (m, 1H, CH). ^13^C NMR (DMSO-*d*_*6*_, δ ppm): 21.16, 21,47, 22.19, 22.43, 22.60, 22.90 (C, CH3, CH3CO); 65.07, 67.70, 69.3, 70.31, 71.3, 73.12 (C, CH); 113.13, 114.63 (C, arom.); 127.5 (C, CH, arom.); 132.14 (C, arom.); 162.63, 162.78 (C, CH); 166.7, 167.1, 167.5, 167.9 (C, CO, CH3CO).

### Animals

The experiments were carried out on adult inbred male CBA mice (n=24), outbred male stock CD1 mice (n=45), outbred female stock CD1 mice (n=40) and outbred male stock CD rats (n=74) (1st category conventional animals). The animals were obtained from the Department of Experimental Biomodelling, E. D. Goldberg Research Institute of Pharmacology and Regenerative Medicine (animal health certificate). Animal maintenance and experimental design were approved by the Bioethics Committee of the E. D. Goldberg Research Institute of Pharmacology and Regenerative Medicine (JACUC protocol No. 96092015) and complied with the directive 2010/63/EU of the European Parliament and the Council of the European Union on the Protection of Animals used for Scientific Purposes, Order No. 199n of the Ministry of Health of Russian Federation (August 1, 2016).

### Study design

When the maximum possible single injection volume of thiowurtzine was used, LD50 (or animal death) was not achieved. The compound was daily administered per os through a probe in a dose range of 50-200 mg/kg (mice) and 25-300 mg/kg (rats) for 1-4 days (Tables 1-3), the last administration was carried out 1 h prior to the pain sensitivity test. The reference drugs were: NSAID diclofenac (Hemofarm), administered orally in a dose of 10 mg/kg in a volume of 0.2 ml/mouse, in a dose of 5 mg/kg in a volume of 0.5 ml/rat, NSAID ketorolac (Dr. Reddy’s Laboratories Ltd..), administered orally in a dose of 10 mg/kg, Tramadol (Organic), administered in a dose of 20 mg/kg via a probe in the form of a solution in purified water in a volume of 0.2 ml/mouse. The suggested doses of reference drugs matched the average therapeutic doses in humans.^43^ The animals of the negative control group received vehicle (water-Tween mixture) in an equivalent volume via the same route.

### Animal methods

The analgesic activity of thiowurtzine was assessed in behavioral tests with thermal and chemical exposures, in Randall-Selitto test with mechanical compression of normal paw pad and with formalin edema.

### Hot plate test

The hot plate test is basic in analgesic activity assessment studies, it involves behavioral responses to pain, controlled by cortical and subcortical structures of brain.^44^ One hour after compound administration, the test was performed using Hot Plate Analgesia Meter (Columbus Instruments). The rats were placed on a plate heated to 54.0±0.5°C, after which pain reaction latency was recorded (licking front and hind paw pads). Analgesic activity was measured by the mean latency in the group and percentage of the pain response inhibition (%PRI), according to the formula: (Tcontrol - Texperiment)/Tcontrol×100%, where T is pain response latency in the corresponding group.

### Acetic acid writhing test

The acetic acid writhing test is aimed at assessment of acute visceral deep pain.^43 45 46 47 44 48^. The specific response to pain (writhing) was caused by intraperitoneal injection of 0.75% acetic acid solution in a volume of 0.1 ml/10 g body weight. The analgesic effect was evaluated by the ability of the compound (within 20 min after the injection) to reduce the number of writhings (in %) in comparison with the control group.

### Randall-Selitto test

The Randall-Selitto test (Randall and Selitto, 1957), intended to serve as a tool to assess the effect of analgesic agents on the response thresholds to mechanical pressure stimulation, usually using by a number of investigators to evaluate inflammatory painful responses.^43 45 46 47 44 48^ An Ugo Basile electronic device based on the Randall-Selitto principle, which allows testing pain in a quantitative manner by pressuring hind limbs of animals, has been used. Hyperalgesia was induced by subcutaneous injection of 2% formalin solution in a volume of 50 μl intraplantarily into the right hind paw pad after one hour compound administration. The Randall-Selitto test was performed twice on each individual one and four hours after compound administration on the left hind paw pad and in first 1-5 min and 40-50 min after the inducing of hyperalgesia on the right hind paw pad. Analgesic activity was evaluated by the ability of the compound to change the response threshold in comparison with the positive and negative control groups.

The rats were sacrificed by CO2 inhalation, the mice were killed by craniocervical dislocation.

### Statistical processing

Statistical processing of the results was performed using ANOVA (Statistica 6.0). For all data, the mean (X) and standard error of the mean (m) were calculated (shown in tables); nis the number of variants in the group. Differences in the studied parameters were assessed with the nonparametric Mann—Whitney U test for testing hypotheses about the homogeneity of the means. The differences were significant at p<0.05.

### Homology Modeling

The mu, kappa, delta and ORL1 opioid receptors were constructed using the corresponding human sequences (respective UniProtKB identifier P35372, P41145, P41143, and P4114) and the online software I-TASSER (Iterative Threading ASSEmbly Refinement) from the Zhang Lab.^40^ This iterative modeling software can use several different partial templates when no single valid template is available. Five models for each receptor were computed using the software default options, and the five best ones were further imported into the MOE software to be checked, charged, and minimized using the AMBER 14:ETH force field.

The homology model for the Ca_v_ 1.2 calcium channel was built using the UniProtKB sequence Q13936 (isoform 1) of the human voltage-gated calcium channel subunit α_1c_ Ca_v_ 1.2 and the PDB structure 6JP5 of the rabbit Ca_v_1.1 / nifedipine complex (resolution 2.90 Å) as a template. The alignment statistics were calculated as follows: 66.4% identity, 75.3% similarity, 4.7% gaps for a total length of 1359 residues. Prior to the homology modeling, the geometry of all the molecules, bonds, and charges of the PDB structure 6JP5 were checked using the Structure Preparation tool of the Molecular Operating Environment software (MOE 2020, Chemical Computing Group, Köln, Germany) and minimized with the Amber 14:EHT forcefield implemented within the MOE software. The homology model was generated using the α_1C_ subunit as a template and the MOE standard parameters in the Protein / Homology Model module. Several loops (215-232, 446-509, 776-892, 941-962, and 1326-1374) were added as the corresponding residues did not exist in the template, but their structure and position were checked and none of them was involved in the interaction with nifedipine.

### Docking

The docking procedures were performed with the GOLD^39^ software (version 2020.3), using the HERMES ^49^ interface (CCDC, Cambridge Crystallographic Data Center) and the implemented GOLD Wizard. We used the GOLD standard parameters for the docking, with the exploration of a spherical site (10 Å radius) centered on the previously co-crystallized ligands. The Genetic algorithm (GA) parameter was set to a number of 200 poses with the activation of the early stop function. We used the ASP (Astex Scoring Potential) fitness function^50^ for calculating the docking scores, and the Chemscore scoring function^41 42^ for the rescore of the results.

### Molecular Dynamics

Molecular Dynamics simulations were performed for the homology model of the Ca_v_ 1.2 calcium channel in presence of thiowurtzine using the Amber 14:EHT force field implemented in the MOE software. A lipid bilayer surrounding the calcium channel and composed of DOPE (1,2-Dioleoyl-sn-glycero-3-phosphoethanolamine) and DOPG (1,2-Dioleoyl-sn-glycero-3-phospho-rac-1-glycerol)-with a respective ratio of 3:1 was generated with the MOE lipid generator, using the PACKMOL-Memgen approach. ^51 52 53^ TIP3P water molecules ^54 55^ were added, as well as Na+ and Clions in order to simulate a 0.1 M ionic force, and pH was set at 7.4. Standard Amber 14:EHT parameters were used for the molecular dynamics. A quick minimization was performed prior to the start of the MD simulations. The geometry of all the molecules, bonds, and charges were then checked using the Structure Preparation tool in MOE, in order to start the molecular simulations. After a preliminary equilibration phase, a 200 ns molecular simulation was performed. A frame was saved every 200 ps, leading to a total number of 1,000 frames.

## ASSOCIATED CONTENT

### Supporting Information

The Supporting Information contains the 1H and 13C NMR spectra for compounds **I, IV, VIII** and **IX**, IR spectra for **I**. It is available free of charge on the ACS Publications website: http://pubs.acs.org…

The structural PDB data for the different models of the receptors (mu, kappa, delta, ORL1, and Ca_v_ 1.2) are also available.

## AUTHOR INFORMATION

### Notes

The authors declare that they have no competing financial interests.

## ACKNOWLEDGMENT

The authors would like to thank Johnny Truong for his work on the homology model of the Ca_v_ 1.2 calcium channel and the subsequent docking of thiowurtzine.

## ABBREVIATIONS

GOLD software: Genetic Optimization for Ligand Docking software
MD: Molecular Dynamics
MOE software: Molecular Operating Environment software
NSAIDs: Non-Steroidal Anti-Inflammatory Drugs
ORL1: Opioid Like Receptor 1
vs: versus

## Notes

### Competing Interest Statement

The authors have declared no competing interest.

